# Redox stress agents strongly enhance mutagenesis during horizontal gene transfer in bacteria and leave distinct mutational and metabolic footprints

**DOI:** 10.64898/2026.06.04.730102

**Authors:** Libertad García-Villada, Brett A. Shore, Kasey Kiser, Isabella G. Russ, Scott A. Gabel, Geoffrey A. Mueller, Natalya P. Degtyareva, Paul W. Doetsch

**Affiliations:** Genome Integrity and Structural Biology Laboratory, National Institute of Environmental Health Sciences, Durham, NC 27709, United States; Nuclear Magnetic Resonance Research Core Facility, National Institute of Environmental Health Sciences, Durham, NC 27709, United States; Research and Development, RTD Biosciences, Charleston, SC 29405, United States; Department of Biological Sciences, North Carolina State University, Raleigh, NC 27607, United States

**Author notes:** Correspondence may also be addressed to Natalya P. Degtyareva.

## Abstract

Redox stress induces DNA mutations that contribute to chronic conditions affecting human health and to the emergence of antibiotic resistance. Yet, the impact of redox stress-induced mutagenesis remains difficult to decipher because redox agents are diverse and produce hard-to-detect mutational outcomes. Single-stranded DNA (ssDNA) provides a useful tool for studying mutagenic effects of redox agents, as it is particularly susceptible to damage and cannot be repaired by most DNA repair pathways. Here, we established a protocol to investigate redox stress-induced mutagenesis based on the *Escherichia coli* conjugative ssDNA that is transferred from donor to recipient cells. Using the environmentally relevant redox agents, potassium bromate and hydrogen peroxide, we show that the F episome is remarkably sensitive to weak mutagens during conjugation, enabling the detection of significant differences in mutational spectra induced by these agents. We support our findings with metabolomic analysis, which reveals agent-specific responses in *E. coli*. We compare these results with those obtained using a yeast ssDNA reporter and conclude that redox-induced mutagenesis depends, among other factors, on the metabolic context of the analysed system. These findings have important implications because the high sensitivity of conjugation-associated ssDNA to environmental mutagens may contribute to the evolution of antibiotic resistance.

**GRAPHICAL ABSTRACT:** 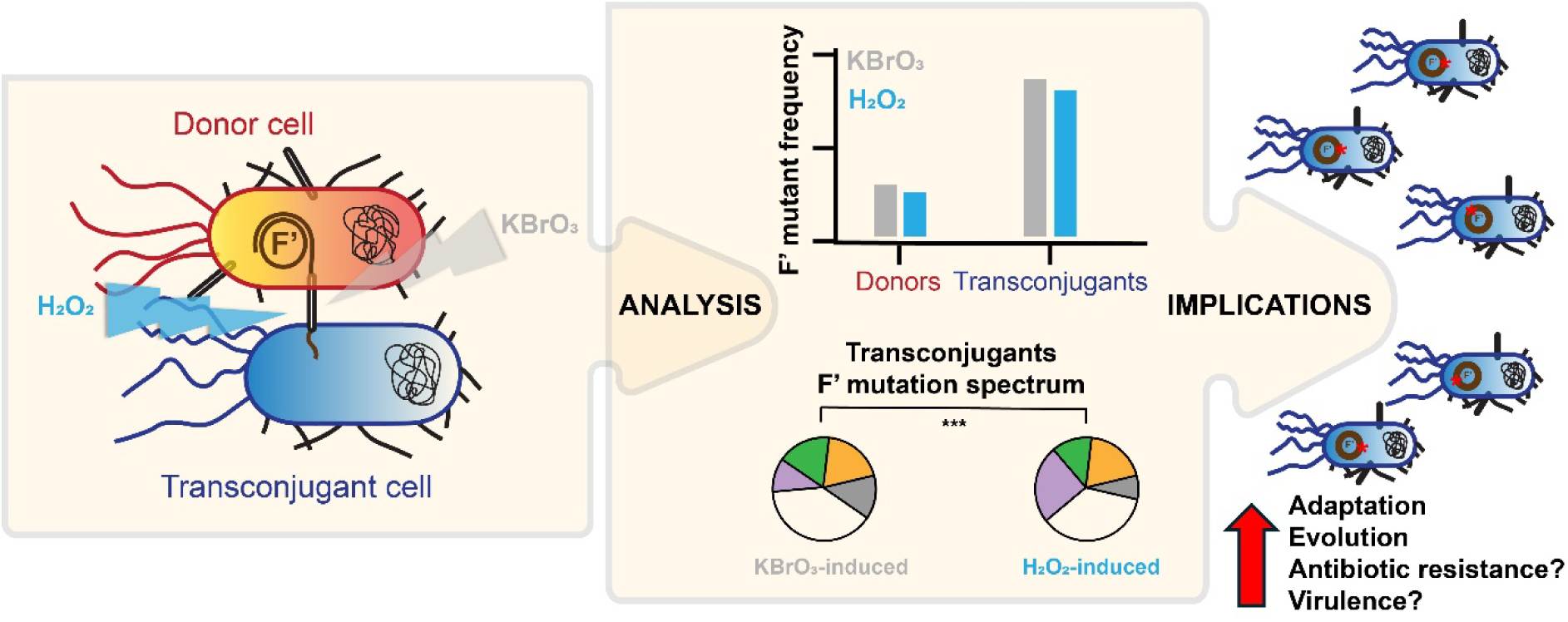

## INTRODUCTION

Redox stress is implicated in the etiology and progression of many chronic conditions that significantly impact human health (reviewed in (1,2), such as cardiovascular and inflammatory diseases (3), neurodegenerative disorders (4–6), and cancer (7–9), as well as aging (10,11). Compared to other known sources of DNA damage, redox stress stands out as a ubiquitous and persistent contributor to mutagenesis (12). Yet, the nature of redox stress-induced mutagenesis is difficult to define, as oxidizing agents are diverse, exert distinct influences on cellular metabolism, and produce subtle, hard-to-detect mutagenic outcomes.

Redox stress manifests from an intracellular imbalance between reactive oxygen species (ROS) and antioxidant defences. ROS are generated as byproducts of normal metabolic processes such as cellular respiration and peroxisomal beta-oxidation, or can be produced during immune cellular respiratory bursts. Atmospheric oxygen and other environmental factors such as ultraviolet and ionizing radiation, cigarette smoke, industrial chemicals, and ozone impose a redox stress threat to normal cellular metabolism (reviewed in (2)). Prokaryotic and eukaryotic cells have evolved several defences against ROS, including enzymatic and non-enzymatic antioxidants, DNA damage signalling and repair mechanisms, and checkpoint control systems (reviewed in (13)). When these defences are surpassed or fail, ROS react with different cell components (fatty acids, proteins, RNA, and DNA) adversely modifying their structure and function.

Previous studies have demonstrated that ROS, through direct oxidative DNA damage (14) or activation of the bacterial RpoS-mediated general stress response (15), are also implicated in the origin and progression of antibiotic resistance (reviewed in (16)). In fact, oxidative stress is one of the main sources of endogenous DNA damage in bacteria not exposed to antibiotics (17). In addition, during infection, bacterial populations are permanently challenged by ROS from multiple sources: ROS produced by host immune cells, which release various oxidizing agents in response to infection, and ROS generated inside of the bacterial cells as a consequence of antibiotic exposure (18).

DNA damage and mutagenesis (spontaneous and induced) in bacteria have been thoroughly studied over time employing the *Escherichia coli* F episome (Fertility factor) as model system (e.g., (19–22)). In this sense, F is particularly useful because it can be maintained either as an independent plasmid or integrated into the chromosome; it can be easily transferred between bacteria; it supports simple genetic manipulation; and it can harbour large amounts of cloned DNA, among other reasons. There is also extensive literature on mechanisms of stress induced mutagenesis based on the F episome (e.g., (23–25)). The F episome, like many other bacterial plasmids, is a conjugative plasmid. It has been well established that conjugative plasmids promote homologous recombination in recipient chromosomes, while also driving episomal rearrangements and, under certain conditions, recombination-dependent mutagenesis on the episome itself (26–30). Yet, little is known about spontaneous or induced DNA damage and mutagenesis of the F episome during conjugation.

Bacterial conjugation is a process of horizontal DNA transfer between a donor cell and a recipient cell by means of direct physical contact through a tube-like structure called pilus (Fig. 1). The involved genetic material is transferred in the form of single-stranded DNA (ssDNA), so that prior to being transferred, both strands of the double-stranded DNA (dsDNA) conjugative molecule are separated, one of them being kept and replicated, to recover its dsDNA structure, in the donor, and the other being transferred to and replicated in the recipient. It thus comprises a system in which long stretches of ssDNA originate and exist for some time until they are copied once the transfer of the genetic material to the recipient cell is completed in approximately 4 minutes (31). ssDNA is known to be particularly susceptible to damage (32), since, unlike the dsDNA, it is not protected in a double helical duplex structure through Watson-Crick base pairing and base stacking. In addition, damage in ssDNA cannot be repaired by DNA excision repair pathways because it lacks the required complementary template strand. The vulnerability of ssDNA to damage presents a unique opportunity of studying the biologically relevant consequences of redox stress under physiological conditions of conjugation.

**Figure 1.**
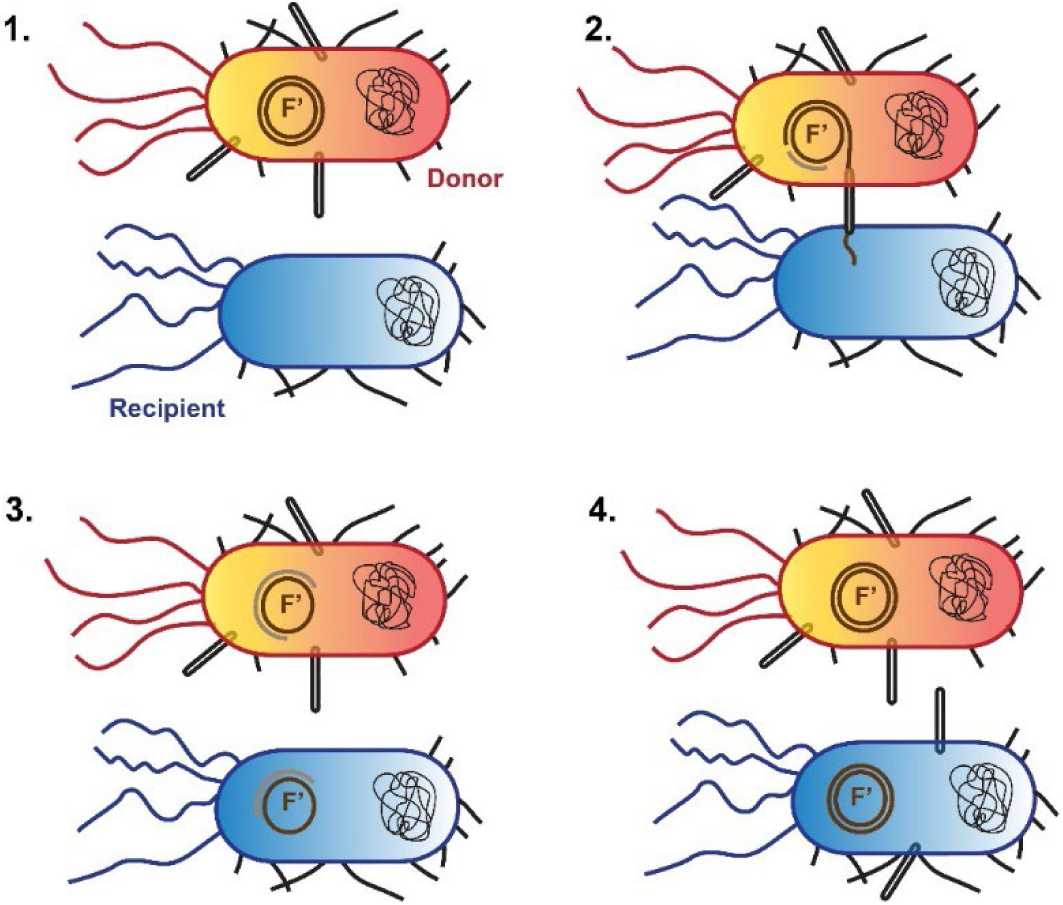
Schematic illustration of bacterial conjugation mediated by the F episome. **1**. The Donor cell carries one or several copies of the F episome, which carries genes encoding the sex pilus. **2**. A sex pilus allows physical contact between a Donor and a Recipient cell. **3**. An F episome in the Donor cell is nicked. This nicked DNA strand is transferred to the Recipient through the sex pilus. F ssDNA in Donor and Recipient are copied to form a dsDNA episome. In the Donor, this process is synchronous with the ssDNA transfer; in the Recipient, it takes place after the DNA transfer is completed. **4**. Recipient cell can now function as a new Donor cell.

Understanding the effect of redox stress on episome mutagenesis during conjugation is critically important considering the key role that conjugative plasmids and conjugation, through horizontal gene transfer, play in the acquisition of new bacterial properties, such as pathogenicity and antibiotic resistance (reviewed in (33–37)). Such findings may be relevant for further delineation of the mechanisms of bacterial adaptation and evolution and potentially can be translated to provide insight into carcinogenesis in vertebrates, since ssDNA is more abundant in many cancer cells compared to normal cells due to replication stress and continuous replication (reviewed in (38)).

In this study, to investigate redox stress-induced mutagenesis during conjugative ssDNA transfer, we took advantage of the method used by Kunz and Glickman (39) to study spontaneous mutagenesis during conjugation: selection of *lacI* mutants. These mutants carry a mutation in the *lacI* gene, which encodes LacI, the lactose operon (*lac* operon) repressor (in our experimental strains, the *lac* operon is located in the F episome). Mutations in this gene that impair repressor function render the lactose operon constitutively activated, and *lacI* mutants can be selected in growth medium where the only carbon source is phenyl-β-D-galactoside (Pgal). Pgal is a lactose analogue that can serve as a carbon source but does not induce the *lac* operon and hence is suitable for detecting constitutive mutants. Mutations in *lacO* (*lac* operator), a short DNA sequence adjacent to *lacI* where LacI binds to repress the *lac* operon, also produce constitutive mutants. The *lacI* system has been extensively used for over five decades and is a well-characterized and reliable model for studying mutagenesis (reviewed in (40–42)).

We employed two different redox stress agents, potassium bromate (KBrO_3_) and hydrogen peroxide (H_2_O_2_). Potassium bromate is used as a food additive in various baked goods as well as other commercial products in the United States (43), while in other regions, such as Europe, Canada and Asia, its use has been banned due to concern for potential health effects, including cancer and reproductive damage (44,45). As the entry point for this compound is the digestive system, it is likely that not only the intestinal cells, but also the intestinal microbiota are exposed to it. The effect of potassium bromate on bacterial populations is currently unknown. On the other hand, hydrogen peroxide is a ROS generated by bacterial cell metabolism, naturally or induced by the action of various antibiotics (46). In addition, it is released by immune system cells during infection (reviewed in (47)). Thus, hydrogen peroxide is a major environmental stressor for bacteria.

Our results show that potassium bromate and hydrogen peroxide significantly increase mutagenesis during conjugation through damage to conjugative ssDNA, resulting in distinct mutational spectra, and induce different metabolic changes that probably underlie their mutational dissimilarities. We also characterize conjugation as a ssDNA-specific mutagenesis system in bacteria to study DNA damage induced by weak mutagens, such as potassium bromate and hydrogen peroxide, and compare the results obtained with this system to the data acquired for the same oxidizing agents with a ssDNA-specific mutagenesis yeast reporter system. We conclude that DNA damage and mutagenesis induced by oxidizing agents depend not only on the specific agent, but also on the specific metabolic characteristics of the system under study. In addition, we consider the potential implications for bacterial evolution and antibiotic resistance for the process of conjugation being highly sensitive to environmental mutagens.

## MATERIAL AND METHODS

### Experimental strains

Unless otherwise indicated, conjugation experiments were performed with the *E. coli* wild-type (WT) strain NR9102 (donor): *ara thi* Δ/*prolac*/ F′/*prolac*/ (F′128-27) (48), and spontaneous nalidixic acid (NAL)-resistant derivatives of the *E. coli* strain S90C (recipient): *ara thi* Δ/*prolac*/ *strA* (48) selected on Lysogenic Broth (LB) agar medium containing 40 µg/ml of NAL (Goldbio). In the experiments performed with the *lacZ* mutation reporter system, a NAL-resistant derivative of the *E. coli lacZ* strains NR10832 and NR10835 (22), with genotype *ara thi* Δ/*prolac*/ and carrying F′CC102 or F′CC105, respectively, were used as donor. The *lacZ* gene present in F′CC102 or F′CC105 contains a base substitution at codon 461 (the original GAG was modified to GGG and GTG, respectively), so that donor and transconjugant cells carrying any of these episomes are unable to grow in a medium in which the sole carbon source is lactose unless the base substitution is reverted (22). All the strains used in the present study were a gift from Dr. Roel Schaaper (NIEHS).

### Conjugation and *lacI* mutant selection

Experiments were conducted as in (39). In brief, spontaneous donor *lacI* mutants were obtained through a series of fluctuation tests (49), in which strain NR9102 was first streaked from a −80°C stock into minimal medium (MM; composition per liter: MgSO_4_(H_2_O)_7_, 0.2 g; K_2_HPO_4_, 10.0 g; NaNH_4_HPO_4_(H_2_O)_4_, 3.5 g; citric acid monohydrate, 2.0 g; thiamine HCl (Thermo Scientific), 5 mg; D-glucose (Gibco), 0.4%; agar (Thermo Scientific) 16 g), lacking L-proline to ensure the presence of the F*’*/*prolac*/ in the growing cells, and incubated. Each experiment started from a single colony grown overnight (ON) in LB medium at 37°C with shaking. The next day, the cell culture was diluted 10^6^-fold in fresh LB, grown for two and a half hours and then distributed in 3-ml aliquots over 20 to 40 independent tubes and incubated ON at 37°C with shaking. The following day, a 50-µl aliquot of each culture was plated on MM agar supplemented with phenyl-β-D-galactoside (750 µg/ml; GBiosciences) as the sole carbon source (Pgal medium) and incubated at 37°C for 48 hours. One mutant colony was collected per plate and streaked on Pgal medium. After this passage, one colony per isolate was independently placed in culture tubes with 3 ml of LB medium. Cultures were grown ON at 37°C with shaking and next day a 0.6 ml aliquot of each was mixed with 0.4 ml of a 50% glycerol solution in a freezing vial and stored at −80°C.

Transconjugant *lacI* mutants were isolated after conjugation performed as follows. Donor and recipient strains were streaked on MM agar (supplemented with L-proline (1 mM; Millipore Sigma), Streptomycin (STR, 200 µg/ml; Goldbio) and NAL (40 µg/ml) for the recipient strain) and incubated. Each conjugation experiment started from a single colony of each strain placed in LB medium (supplemented with STR and NAL for the recipient strain) and grown ON at 37°C with shaking. Next day, donor and recipient cultures were diluted 100-fold in fresh LB medium and grown for about two additional hours till OD_600_ reached ∼0.5. Then cultures were collected, concentrated 10-fold, washed three times with M9 salts (composition per liter: KH_2_PO_4_, 3.0 g; Na_2_HPO_4_(H_2_O)_7_, 12.8 g; NaCl, 0.5 g; NH_4_Cl, 1.0 g; Gibco), resuspended in M9 medium (same composition per liter as M9 salts but also contains: MgSO_4_, 241.0 mg; CaCl2, 11.1 mg, D-glucose, 0.4%), and mixed (in a ∼1:1 donor:recipient proportion). In experiments where cells were exposed to a mutagen during conjugation, the mixed culture was divided in two identical volumes (of 20 ml each): one was used as control (mock), and the other was exposed to a concentration of mutagen (1.5 mM for potassium bromate, KBrO_3_; 10 mM for hydrogen peroxide, H_2_O_2_; and 10 mM for methyl methane sulfonate, MMS, unless otherwise indicated; Millipore Sigma). Cultures were incubated at 37°C without shaking. Conjugation was stopped after one hour by adding NAL to a final concentration of 40 µg/ml, and by vigorously vortexing the cultures and placing them on ice. After 15 minutes, cultures were washed once with, and resuspended in, M9 salts. A sample of each culture (mock and exposed) was serially diluted in M9 salts and plated on: (i) MM agar supplemented with L-proline, STR and NAL, to determine total recipient cell density (pro^−^ NAL^R^ STR^R^ selection) and (ii) MM agar supplemented with STR and NAL (but not L-proline), to determine total transconjugant cell density (pro^+^ NAL^R^ STR^R^ selection). Samples from both cultures (mock and exposed) were also directly plated on Pgal medium supplemented with STR and NAL to determine the cell density of transconjugant *lacI* mutants (*lacI* pro^+^ NAL^R^ STR^R^ selection). A sample of the donor strain was retrieved before conjugation and plated on (i) MM agar (after serial dilution in M9 salts) and (ii) Pgal medium, to determine its total donor cell density and *lacI* mutant density, respectively. Plates were incubated at 37°C and colonies were scored after 28 or 48 hours for MM agar and Pgal medium, respectively. *lacI* mutant frequencies were calculated as the ratio between *lacI* mutant density per ml and total transconjugants per ml. The possibility of NAL inducing mutagenesis on the selective plates was ruled out by control experiments in which 50-µl aliquots from ON *lacI*+ transconjugant cultures were plated on Pgal medium without (mock) and with NAL (40 µg/ml) and the frequency of *lacI* colony forming units (CFU) was compared between both types of Pgal selective plates. All media used was freshly made for each individual experiment.

### Non-conjugation experiments

Non-conjugation control experiments were conducted as the conjugation ones described above but only with the donor strain.

### *lacZ+* mutant selection

For these experiments, donor strains were NAL-resistant derivatives of the *lacZ* strains NR10832 and NR10835 (described above) and the recipient strain was S90C. Conjugation was conducted as detailed above for the *lacI* reporter with a few modifications: (i) for mutant isolation, lactose (0.4%; Millipore Sigma) medium was used instead of Pgal medium; and (ii) donor and transconjugant *lacZ+* revertants were isolated and their frequencies determined (NAL and STR were used to select for donor and transconjugant *lacZ+* mutants, respectively).

### *rpoB* mutant selection

Experiments were conducted similarly to the non-conjugation experiments (above) with the following differences. After exposure, cells were washed once with M9 salts and then resuspended in LB medium and grown for 20 hours at 37°C with shaking before 0.2 ml samples were retrieved and plated on LB agar supplemented with rifampicin (RIF, 100 µg/ml; Goldbio). Survivors were determined by plating diluted samples of the experimental cultures on LB agar immediately after exposure and after 20 hours of recovery. *rpoB* mutant frequency was estimated as the ratio between rifampicin-resistant (RIF^R^) mutant density per ml and survivors at 20 hours of recovery per ml.

### Metabolite extraction

For metabolite extraction, cells (donor strain NR9102) were handled as in the non-conjugation experiments, but the cultures’ volume was doubled to get enough cells for the analyses. After exposure to KBrO_3_ (1.5 mM) or H_2_O_2_ (10 mM), cells were harvested and washed one time with M9 salts. A sample of the cultures (treated and mock) was serially diluted and plated on MM agar to determine the percentage of survivors. Cultures were then washed one time with PBS (pH 7.4; Gibco), resuspended in 1 ml PBS, and frozen at −80°C. For the analysis, samples were thawed on ice and sonicated 20 seconds at 30% power using a micro-sonicator unit (Q Sonica). Solutions were centrifuged 15 k*g* for 20 minutes at 4°C. The supernatant was passed through a 10 kDa-cutoff concentrator that was prerinsed to remove residual glycerol. The flow through was frozen and lyophilized. Dried samples were dissolved in 550 ml of ^2^H-phosphate buffered saline with 0.25 mM azide and 0.1 mM DSS (3-(trimethylsilyl)-1-propanesulfonate, 2,2-dimethyl-2-silapentane-5-sulfonate; Cambridge Isotope Laboratories) for concentration and chemical shift reference.

### 1H-NMR spectroscopy

NMR spectra were acquired on an 800 MHz Agilent DD2 console equipped with a cryogenically cooled probe. A standard 1D-NOESY sequence with 4 seconds acquisition time, 1 second recycle delay, 256 transients, and 100 milliseconds mixing time was utilized.

### Metabolomics data analysis

For each sample, the metabolite concentrations determined from the ^1^H-NMR spectrum were square-root transformed and then auto scaled (mean-centred and divided by the standard deviation of each variable). Chenomx software (Alberta, Canada) was used for processing and spectral analysis. Data normalization and analyses were performed using MetaboAnalyst 6.0 software.

### Sanger sequencing

*lacI* colonies were isolated from the conjugation selective plates and grown ON in LB medium to create frozen stocks. A sample from these cultures was used to amplify the *lacI* through PCR (for information on the primers, see Supplementary File S1), and then the amplification products were Sanger sequenced (Genewiz) and analysed for mutations using CLC Genomics Workbench 23.

### Library preparation and whole-genome sequencing of the F episome

To verify the integrity of the F episome carried by donor strain NR9102 and determine the *lacI* DNA strand that is transferred from donor to recipient during conjugation, we sequenced the F from this strain (F′128-27) (48). We also sequenced the episome of a small sample of *lacI* mutants from conjugation experiments conducted with MMS (10 mutants from the mock population and 10 mutants from the exposed population) looking for additional mutations at F. In all cases, the episome DNA was extracted using a Miniprep Kit (Promega) and its quality determined in an Agilent 2100 TapeStation. Libraries were prepared with Celero^TM^ EZ DNA-Seq Kit (Tecan) and sequenced with NextSeq 550 System Mid-Output Kit (Illumina Inc, USA) in a NextSeq 500 System Whole-Genome Sequencing Solution (Illumina Inc, USA). The resulting paired-end reads were 150 bp long.

### Variant calling

CLC Genomics Workbench 20 was employed as analytical pipeline. Reads were first trimmed and then aligned to the F reference genome (GenBank CP014271). Once aligned, removal of duplicates and local realignment of reads around InDels were performed with default settings. For variant calling, the minimum central quality was 30, the minimum neighbourhood quality was 20, and two minimum variant allele frequency (VAF) threshold of 35 was used. For InDels and structural variants, the P-value threshold was 0.0001 and the minimum number of reads, 2.

### Statistical analyses

Unless otherwise indicated, all data was analysed using GraphPad Prism 10. The statistical tests used for specific experiments are described in Results and Figure Legends.

## RESULTS

### Redox stress mutagens, potassium bromate and hydrogen peroxide, robustly increase mutagenesis in the F episome during conjugation

To assess induced mutagenesis in the bacterial F episome during conjugative ssDNA transfer between donor and recipient cells, we utilized the *lacI* reporter system. When *E. coli* cells were exposed during conjugation to equitoxic (producing equal toxicity or having equivalent detrimental effects on culture growth under the conditions being compared) concentrations of potassium bromate (1.5 mM) and hydrogen peroxide (10 mM), the frequency of *lacI* mutants increased ∼10-fold in the exposed transconjugant populations compared to mock-exposed cultures (Fig. 2A and B, compare “Conjugation” and “Non-conjugation”). Exposure concentrations were moderately toxic, resulting in death of 85-95% of cells (Fig. 2D). Potassium bromate and hydrogen peroxide are considered “weak” mutagens because, at equitoxic doses (resulting in similar cell survival), they induce fewer mutations than “strong mutagens” such as UV light, for example (50,51). Without conjugation taking place, none of these compounds significantly increased *lacI* mutagenesis (Fig. 2A and B, “Non-conjugation”). Altogether, these data indicate that the redox stress agents potassium bromate and hydrogen peroxide induce mutagenesis during conjugation through damage to F ssDNA. These results were further corroborated by quantification of *lacI* mutant frequencies following exposure to the S(N)2-type alkylating agent methyl methane sulfonate (MMS) during conjugation. ssDNA is considerably more vulnerable than dsDNA to damage induced by MMS (52). As expected, exposure to MMS was hyper mutagenic during conjugation (Fig. 2C). To further verify that potassium bromate and hydrogen peroxide, at the concentrations used in these experiments, do not significantly increase mutagenesis in dsDNA, and therefore the mutant frequency increase mainly results from ssDNA damage, we assessed the induction of rifampicin-resistant (RIF^R^) mutants. RIF^R^ cells arise from mutations in the chromosomal gene *rpoB*. Since our donor strain does not have the F episome integrated into the chromosome, its chromosomal DNA is not transferred during conjugation, and a chromosomal gene may serve as reporter for mutagenesis in dsDNA. We did not observe significant induction of mutagenesis in chromosomal DNA of exposed donor cells (Supplementary Fig. S1).

**Figure 2.**
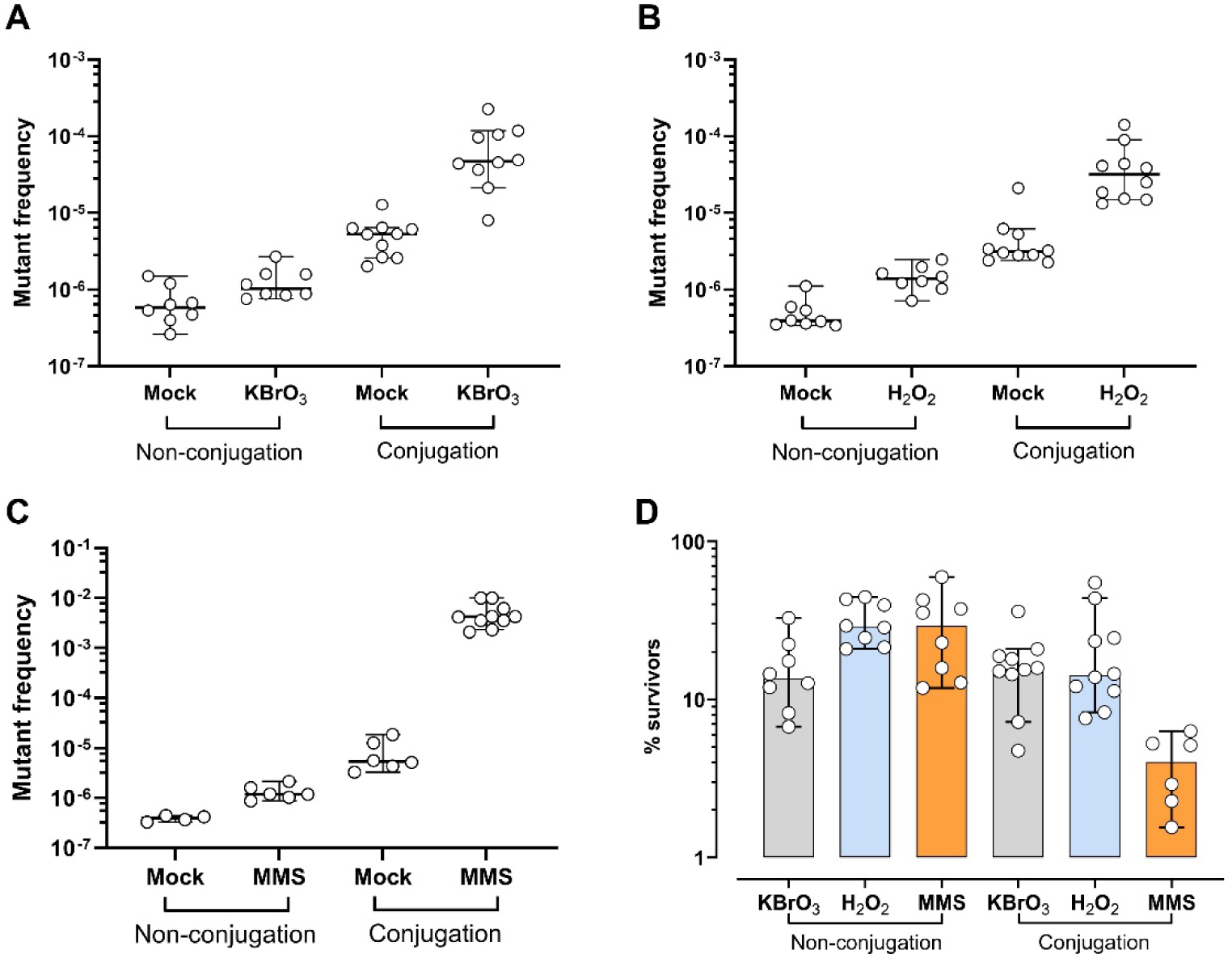
Exogenous stress increases the frequency of mutations during conjugation. Exposure to (**A**) potassium bromate (1.5 mM), (**B**) hydrogen peroxide (10 mM), and (**C**) MMS (10 mM) significantly increased the frequency of *lacI* mutants (colony forming units (CFU) of *lacI* mutants per ml / CFU of survivors per ml) during conjugation. In contrast, no significant induction of mutagenesis was observed after exposure in the absence of conjugation (“Non-conjugation” in the figures). (**D**) Percentage of survivors ((CFU of exposed cultures per ml / CFU of mock-exposed cultures per ml) x 100) after exposure to potassium bromate, hydrogen peroxide, and MMS in (**A**), (**B**), and (**C**), respectively. In all figures, the horizontal line and error bars represent median +/− 95% interval of confidence limits of n≥ 6 biological replicates and/or independent experiments. Figure showing the effect of exogenous stress on mutagenesis during bacterial conjugation. Panels A–C present mutation frequencies of *lacI* mutants after exposure to potassium bromate (1.5 mM), hydrogen peroxide (10 mM), or MMS (10 mM). In all three cases, exposure during conjugation significantly increased mutation frequency in transferred DNA compared with unexposed controls. In contrast, exposure under non-conjugation conditions did not significantly increase mutagenesis. Panel D shows survival percentages after exposure to each mutagen, calculated relative to mock-exposed cultures. Data are displayed with horizontal lines representing medians and error bars representing 95% confidence intervals from at least six biological replicates and/or independent experiments.

Conjugation-dependent induction of mutagenesis in the F episome has been previously reported (39,53). However, to our knowledge, the consequences of direct exposure to exogenous damage while conjugation is taking place have not been studied, with a few exceptions (53,54). In contrast to previously published methods for analysis of mutagenesis in ssDNA, in which elegant and complex, but somewhat artificial reporters were constructed in mutant backgrounds, allowing the generation of long stretches of ssDNA (32,55), we have developed a protocol to study mutagenesis in ssDNA during the physiological process of conjugation. In the course of this process, the F episome DNA is extremely vulnerable to damage because it is being transferred from one cell (donor) to another (recipient) through the conjugational bridge as a single-stranded molecule, so it is not protected, like the dsDNA, in a double helical duplex structure through Watson-Crick base pairing and base stacking, or like other ssDNA formations by single-stranded binding proteins (SSBs) (31).

This system proved to be sensitive and robust as a tool for studying the molecular mechanisms of ssDNA mutagenesis for the following reasons. Firstly, under the experimental conditions used in this study, the *lacI* mutant frequency in the transconjugant population after conjugation in minimal (M9) and rich (LB) media was almost an order magnitude higher than in the donor culture before conjugation (Supplementary Fig. S2A), providing a necessary source for statistically meaningful mutational spectra analysis. To verify that this increase was not an artefact resulting from plating the experimental cultures on selective media containing nalidixic acid (NAL), which is a known bacterial mutagen (56), we conducted control experiments in which WT transconjugants (pro^+^ NAL^R^ STR^R^) were plated on Pgal medium with and without 40 µg/ml NAL. The frequency of *lacI* mutants was the same in both selective media (Supplementary Fig. S3). Thus, the observed increase in mutant frequency is consistent with the results of previous studies by Kunz and Glickman (39), which showed that conjugation may increase the total *lacI* mutant frequency by about two-fold when transfer occurs in rich medium (LB). They also found, through the analysis of a mutation spectrum based on 36 characterized amber sites, that it primarily produces base substitutions, mainly G:C to A:T transitions. To our knowledge, no further studies of spontaneous mutagenesis during conjugation have been published. We extended the characterization of the spontaneous mutational spectra of conjugation (Supplementary Fig. S2 and Supplementary Files S1 and S2).

Secondly, we report for the first time the differences in the mutational spectra during conjugation in minimal versus rich medium (Supplementary Fig. S2B). As Kunz and Glickman (39) did, we found, in rich medium, a significant increase in base substitutions in the transferred F population due mostly to an increase in the frequency of mutations at C/G, in particular G:C to A:T, but also G:C to T:A (Supplementary Fig. S2B and C and Supplementary File S2). Since we know (through whole genome sequencing and information published in (57,58)) which DNA strand from the F episome is transferred during conjugation, we can ascribe the G:C to A:T mutations to C to T. Interestingly, of the 45 C to T transitions in mutants selected from rich media, 28 were located in two hot spots: at positions 104 (17 mutations), and at 419 (11 mutations) (Supplementary Fig. S2D, “Post-conjugation in RM”). These two C to T hot spots were found within the same DNA context: cCagg, which are two of the three sequences (there is an additional cCagg at position 959) in *lacI* that represent a specific substrate of Dcm (DNA cytosine methyltransferase). 5-methylcytosine is well known to be more prone in bacteria to cytosine deamination (59). Overall, we do not observe the same changes when conjugation occurs in minimal medium (Supplementary Fig. S2B and C), indicating that cytosine deamination during conjugation may happen more frequently in rich medium. During experimentation, cells are concentrated ten-fold to favour conjugation. In rich medium, this may lead to rapid alkalinization of the environment, since the primary carbon sources in rich medium (LB) are amino acids and peptides whose catabolism generates alkaline byproducts. Alkaline conditions are known to significantly accelerate cytosine deamination (60). These results indicate that the type of growth medium is an important factor to consider when working with sensitive mutation reporters.

This comprehensive and nuanced characterization of F mutagenesis associated with conjugation, in comparison with previously published data, reveals distinct spectra across different media and further attests to the sensitivity of our experimental system. Overall, the data presented demonstrate that conjugation is a highly sensitive system for detecting mutagenesis induced by weak mutagens, including potassium bromate and hydrogen peroxide. Furthermore, most of the mutations generated during conjugation result from damage to the ssDNA that is transferred from the donor to the recipient cell. This makes it possible to identify which specific DNA bases are more susceptible to damage or mutation by particular mutagens, thereby enabling detailed comparisons of the mutational spectra they generate.

### Exposure to potassium bromate or hydrogen peroxide generates distinct mutational spectra

Following mock, potassium bromate, hydrogen peroxide, and MMS exposures, we Sanger sequenced *lacI* and adjacent *lacO* loci in selected transconjugants (Material and Methods). Hydrogen peroxide and potassium bromate are widely used as experimental oxidants to induce redox stress and oxidative DNA damage (reviewed in (61,62)). However, the mutational spectra reflecting exposure to these two agents were significantly different from each other (Fig. 3A). In both spectra, as well as in the mock-exposure spectrum (Fig. 3A, “Mock”), base substitutions comprised a smaller fraction than other types of mutations. The “Other” category of mutations (Fig. 3A and B) comprises: +/− 1 base pair changes (+/− 1 BP), >1 base pair insertions and deletions (InDels), duplications and complex mutations, as well as mutants in which no mutations in the entire *lacI-lacO* region were detected. This category also includes some that could not be amplified by PCR (designated as “null”), presumably due to deletions affecting the amplification primers’ binding site (Fig. 3D). This type of spectrum is also characteristic of spontaneous mutagenesis in the F episome during vegetative replication (Supplementary Fig. S2B, “Pre-conjugation”) and contrasts sharply with the spectrum induced by the strong mutagen MMS, for which only 22% of mutations were not base substitutions (Fig. 3B). The contribution of each sub-type of non-substitution changes is significantly different in mutants selected following exposure to potassium bromate compared to the mutants derived from mock-exposed cells (Fig. 3D). Potassium bromate increased the fraction of +/− 1 BP, specifically −C, −T, and +C mutations (18, 19, and 45 relative fold increases, respectively, when compared to mock; Fig. 3E). The proportion of non-substitution changes induced by hydrogen peroxide was not significantly different from mock exposure (Fig. 3D), but it still showed an 18-fold increase in deletions of a single cytosine relative to mock (Fig. 3E). In most cases, these single base pair insertions and deletions were observed in homonucleotide runs (Supplementary Table S1), which suggests that DNA polymerase slippage was involved in the process of mutagenesis (63). With respect to base substitutions, there is an increase in the relative frequency of cytosine changes in the presence of potassium bromate compared to mock (Fig. 3C), while hydrogen peroxide induced an increase in the relative frequency of cytosine and guanine changes (Fig. 3C). There is a prevalence for C to A mutations in the spectrum induced by potassium bromate, and for C to T, G to A, and G to T mutations in the spectrum induced by hydrogen peroxide (Fig. 3C). Differences between these two agents in the relative frequencies of mutations they induce are not due to the presence of hot spots (Supplementary Fig. S4). In summary, these results indicate that potassium bromate and hydrogen peroxide exposure lead to the mutagenesis of different bases in ssDNA and, thus, that their mechanisms of mutagenesis differ between them.

**Figure 3.**
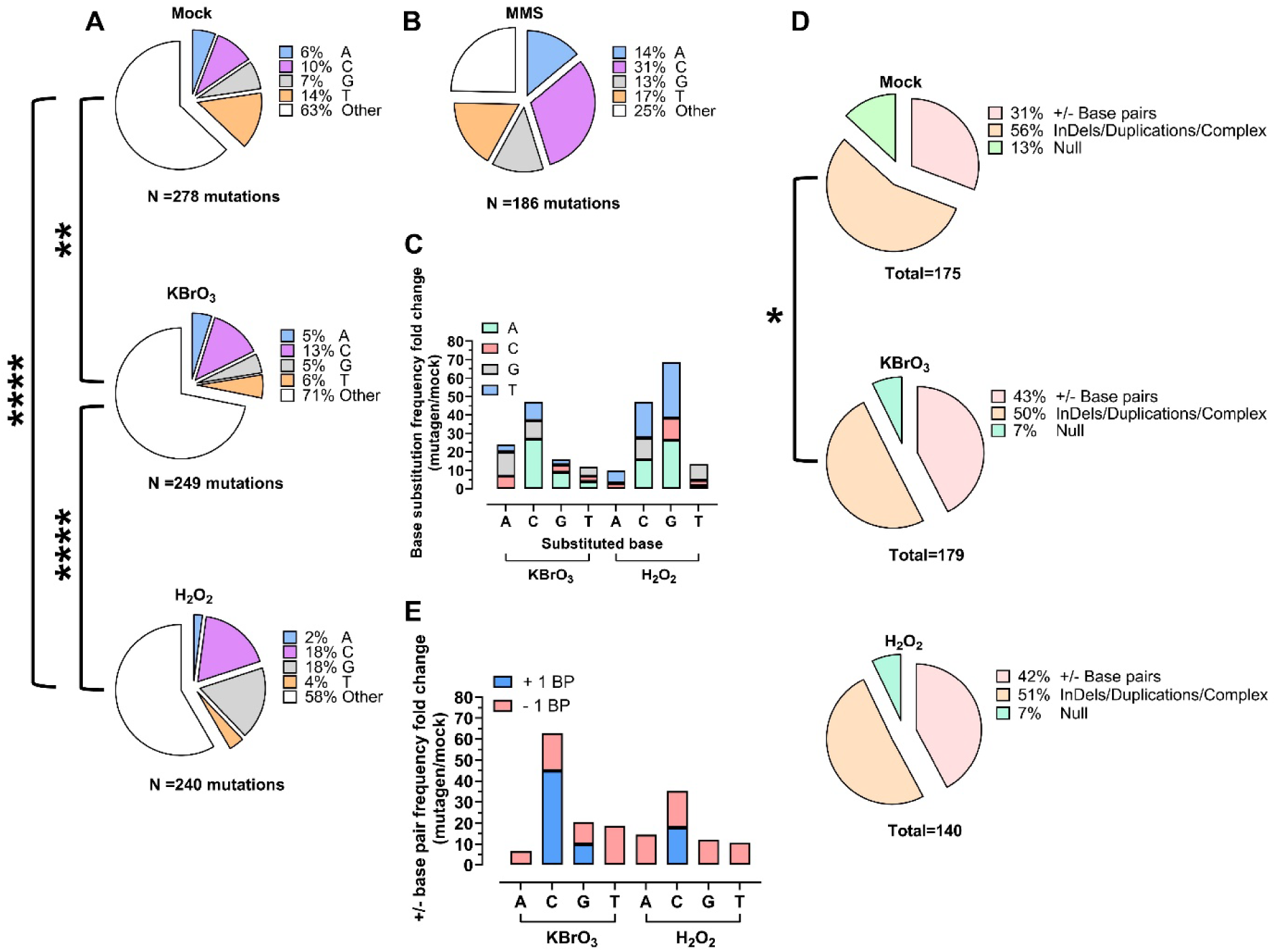
Redox stress agents, potassium bromate and hydrogen peroxide, render a different mutational spectrum during conjugation. (**A**) Spectra of mutations identified by Sanger sequencing of *lacI* mutants from mock-, potassium bromate-, and hydrogen peroxide-exposed transconjugant populations. The overall proportion of each type of mutation in cultures exposed to the oxidizing agents is significantly different from the mutation distribution of mock-exposed cultures (P= 0.0055 and P< 0.0001 for potassium bromate and hydrogen peroxide, respectively; chi-square test). Proportions of mutations were also significantly different between potassium bromate- and hydrogen peroxide-exposed transconjugants (P< 0.0001; chi-squared test). While potassium bromate-exposed mutants show an increased proportion of Other mutations, base substitutions, mainly at C and at G, are prevalent in hydrogen peroxide-exposed ones. (**B**) Spectra of mutations of *lacI* mutants from MMS-exposed transconjugant populations. The proportion of Other mutations is significantly reduced in pro of a major increase in base substitutions. (**C**) Relative fold change in base substitution frequencies of potassium bromate- and hydrogen peroxide-exposed mutants compared to mock-exposed. Potassium bromate induced increased C to A transversions while hydrogen peroxide induced increased mutations at C and at G. (**D**) Distribution of Other mutations in mock-, potassium bromate-, and hydrogen peroxide-exposed transconjugant populations. Potassium bromate-exposed populations show a significant increase in +/− 1 base pair when compared to mock-exposed populations (P= 0.0334; chi-square test). (**E**) Relative fold change in +/− 1 base pair in potassium bromate- and hydrogen peroxide-exposed transconjugants compared to mock-exposed. Potassium bromate induced an increase of mutations at C, mainly +C. Figure showing mutational spectra of *lacI* mutants resulting from bacteria exposed to oxidative stress agents and MMS during conjugation. Panel A compares mutation spectra in mock-, potassium bromate-, and hydrogen peroxide-exposed transconjugant populations determined by Sanger sequencing. Mutation distributions differ significantly between oxidant-exposed and mock-treated samples, and also between potassium bromate and hydrogen peroxide treatments. Potassium bromate exposure increases the proportion of “Other” mutations, whereas hydrogen peroxide exposure predominantly increases base substitutions, especially mutations at cytosine and guanine residues. Panel B shows mutation spectra after MMS exposure, with a marked increase in base substitutions and a reduction in “Other” mutation types compared with mock-treated controls. Panel C presents relative fold changes in base substitution frequencies compared with mock-treated populations. Potassium bromate preferentially induces C-to-A transversions, while hydrogen peroxide increases mutations at cytosine and guanine bases. Panel D shows the distribution of “Other” mutations, with potassium bromate significantly increasing single-base insertions or deletions of one base pair compared with mock-treated populations. Panel E presents relative fold changes in single-base insertions and deletions compared with mock-treated controls, showing that potassium bromate mainly increases mutations at cytosine residues, particularly single-base cytosine insertions.

### Mutagenesis in the donor cells’ F DNA does not significantly contribute to increase mutagenesis during conjugation

During conjugation, not only the DNA strand that is transferred (T-strand) from donor to recipient, but also a section of F episome in the donor cell complementary to the T-strand (the N-strand) exists as ssDNA. A recently published study on the dynamics of conjugative plasmid transfer (31) shows that there is a delay of about 3 minutes between the generation of the N-strand and its duplication in the donor cell, and that the same conjugative episome can be transferred more than once during a single donor-recipient conjugative interaction. Accordingly, the N-strand might also be vulnerable to damage, and a mutation generated during consequent DNA replication can be transferred to the recipient cell in a second round of conjugation. To address the potential contribution to mutagenesis during conjugation from damage to the donor N-strand, we established a system based on a NAL-resistant derivative of the donor strain (NR9102), and a streptomycin (STR)-resistant, NAL-sensitive derivative of the recipient strain (S90C), allowing selection for *lacI* mutations induced in the donor cells during conjugation. We conducted several experiments with this system using hydrogen peroxide (10 mM) and MMS (10 mM) as mutagens. Yet, no significant induction was observed in the donor population (Supplementary Fig. S5).

We conjectured that detection of *lacI* mutants in the donor population after conjugation could be obscured by the presence of more than one F episome in the cell (64), and that, therefore, only dominant mutations would generate a selectable phenotype. To avoid this obstacle, we used a dominant system based on the *lacZ* reporter. *lacZ* encodes the enzyme β-galactosidase, which is a component of the *lac* operon, responsible for hydrolysing lactose into glucose and galactose. Cells carrying this reporter in the F episome are unable to grow in medium in which the sole carbon source is lactose due to a missense mutation in *lacZ* (Material and Methods). However, *lacZ*^+^ revertants, generated through G/C to A/T or T/A to A/T substitutions in our experimental strains, can be selected in lactose medium. Since MMS is a potent ssDNA mutagen, we employed it in initial, proof of principle experiments. In a series of conjugation experiments utilizing the F *lacZ* reporter system, we observed between a 30- and a 40-fold induction of mutagenesis in the donor strain (Fig. 4, “Conjugation”). To exclude the possibility that such an increase is driven by previously unrecognized induction of mutagenesis in dsDNA, we performed the same experiments under non-conjugation conditions. An induction of mutagenesis comparable to what we observed during conjugation occurred in the absence of conjugation (Fig. 4A and B “Non-conjugation”). These results indicate that the induction of mutagenesis observed in the donor in these experiments is not linked to conjugation. Although a 30- to 40-fold induction of mutagenesis is significant, it is relatively minor when compared to the intensity of induction observed in MMS-exposed transconjugants (at least 1,000-fold; Fig. 4)). Collectively, these results, similar to those shown in the previous section, indicate that accumulation of DNA damage at F during conjugation occurs mainly in the T-strand. Such biased, non-symmetrical mutational pressure may have important implications for evolution of antibiotic resistance and virulence under exogenous stress.

**Figure 4.**
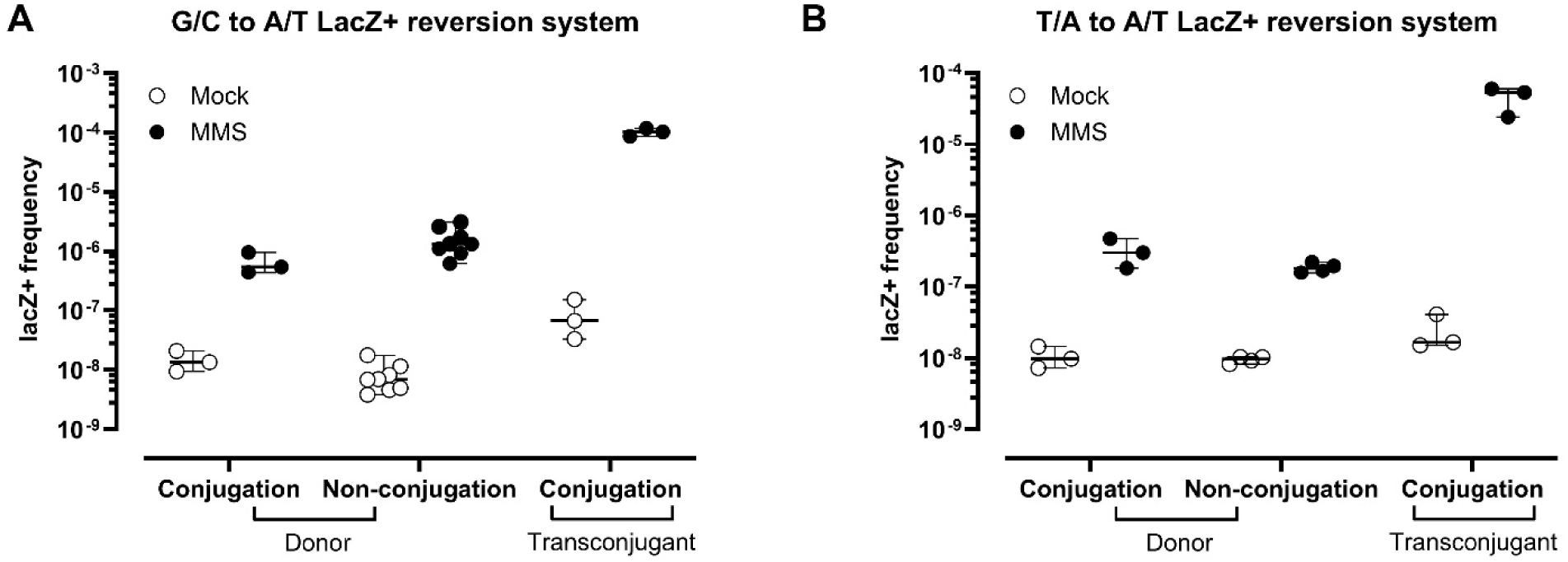
Mutagenesis in the donor cells’ F DNA does not significantly contribute to increased mutagenesis during conjugation. Exposure to MMS (10 mM) during conjugation significantly increased the *lacZ* revertant frequency (CFU of *lacZ* revertants per ml / CFU of survivors per ml) in transconjugant and donor populations in two different *lacZ* reversion systems. The increase in the donor populations was similar under conjugation and non-conjugation conditions, which implies that a fraction of the mutations induced in the F episome during conjugation (13% maximum) is not necessarily linked to conjugation itself. The horizontal line and error bars represent median +/− 95% interval of confidence limits of n≥ 3 biological replicates and/or independent experiments. Figure showing the contribution of donor-cell F episome mutagenesis to mutation frequencies observed during conjugation after MMS exposure. Exposure to MMS (10 mM) increased lacZ reversion frequencies in both transconjugant and donor populations in two independent lacZ reversion assays. Transconjugants displayed elevated mutation frequencies during conjugation, while donor populations showed similar increases under both conjugation and non-conjugation conditions. These results indicate that only a minor fraction of the mutations detected in the F episome during conjugation, estimated at a maximum of 13%, can be attributed specifically to conjugation-associated processes. Horizontal lines represent medians and error bars indicate 95% confidence intervals from at least three biological replicates and/or independent experiments.

### Potassium bromate or hydrogen peroxide cause distinct changes in bacterial metabolism

Significant differences in potassium bromate- and hydrogen peroxide-induced mutational spectra indicate that exposure to these redox stress agents results in distinct mechanisms of mutagenesis and suggest that they might have different, specific effects on cellular metabolism. To compare the consequences of exposure to these agents, we analysed the metabolic profiles of exposed donor populations by NMR spectroscopy. For each oxidizing agent experiments were carried out following the same steps as in the non-conjugation experiments (Material and Methods). By employing the metabolite reference library in Chenomx, we identified and measured the concentration of 34 and 35 distinct compounds in cell extracts from experiments conducted with hydrogen peroxide and potassium bromate, respectively. Principal component analysis (PCA) revealed that the metabolic profiles of cultures exposed to either potassium bromate or hydrogen peroxide were significantly different from the profiles of the corresponding mock cultures (Fig. 5A and B). PC1 and PC2 combined explain ∼60% of the observed variance in both analyses. When all data was merged, metabolic profiles were significantly different from the combined mock population and from each other (Fig. 5C). In this analysis, PC2 separated the populations based on treatment, showing the highest loadings for 4-aminobutyrate (−0.30), glycine (0.27), isoleucine (0.22), N-acetylputrescine (−0.30), NADP+ (−0.24), sucrose (0.28), valine (0.30), and dTTP (−0.23) (Supplementary File S3). To normalize the difference observed between potassium bromate- and hydrogen peroxide-exposed cultures, we determined and compared the fold change in concentration (relative to the corresponding mock cultures) for each of the identified metabolites in the experimental populations (Fig. 5D). For all comparable metabolites, there was no significant difference in the average fold change between potassium bromate- and hydrogen peroxide-exposed cultures (paired t test, P= 0.9185; data normalized using square root transformation). Yet, for several individual metabolites, the 95% interval of confidence for fold change differences do not overlap between treatments, indicating the existence of a significant difference in their concentration per cell (Fig. 5D). These metabolites are glutathione, methionine, N-acetylputrescine, 4-aminobutyrate, and uracil. The first two display lower concentrations following potassium bromate exposure, and the last two display higher concentrations after hydrogen peroxide exposure; N- acetylputrescine also displays higher concentration after potassium bromate exposure. Although these compounds are mapped in different metabolic pathways, we can suggest a plausible reason for changes in their levels due to oxidative stress. For example, exposure to potassium bromate specifically produces depletion of cellular glutathione (Fig. 5D and (51)). This interaction may also explain the greater decrease in the levels of methionine in cells exposed to potassium bromate, since methionine is used to produce cysteine, which then serves as a building block precursor for glutathione. N-acetylputrescine, the N-acetylated form of the antioxidant putrescine, is linked to polyamine metabolism, and its concentration often increases during redox stress as a product of putrescine metabolism (65), and 4-aminobutyrate (GABA) is also an intermediate in putrescine degradation. Uracil is a product of cytosine deamination, and its levels are increased in both potassium bromate- and hydrogen peroxide-exposed cells, but such increase is most pronounced in the hydrogen peroxide-exposed ones, which also mirrors the increased percentage of C to T mutations observed in hydrogen peroxide exposed cells (Fig. 3C).

**Figure 5.**
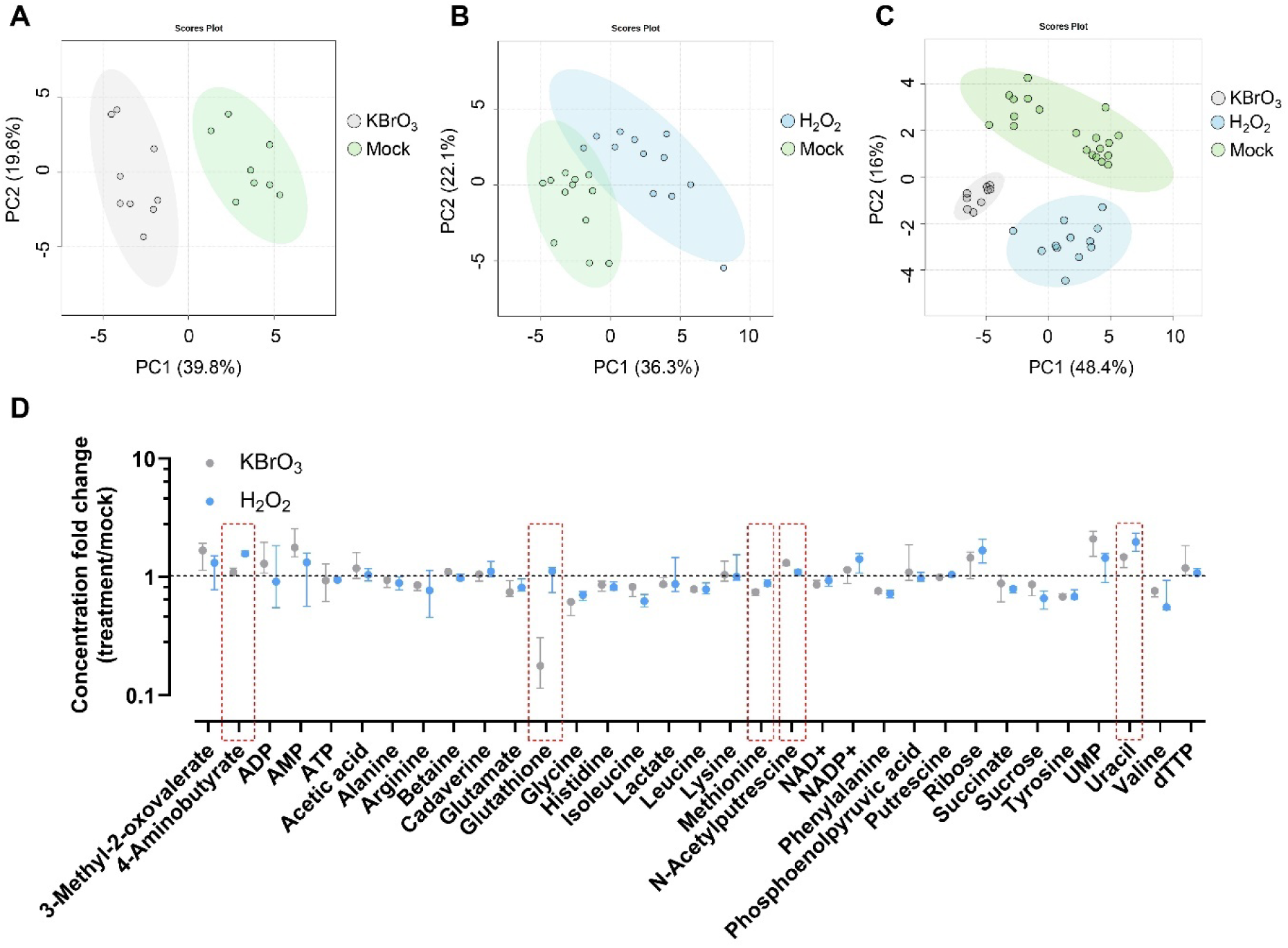
Potassium bromate and hydrogen peroxide cause distinct changes in bacterial metabolism. Principal component analysis (PCA) of NMR-detected metabolites in donor cells exposed to (**A**) potassium bromate (1.5 mM) and (**B**) hydrogen peroxide (10 mM) reveals significant differences in the metabolic landscapes of stress- and mock-exposed populations. (**C**) The metabolic landscapes of populations exposed to one or another oxidative agent were also significantly different. In (A), (B), and (C), each dot represents one of three biological replicates from a total of three (potassium bromate) or four (hydrogen peroxide) independent experiments. Coloured ovals represent 95% confidence intervals. PC, principal component. (**D**) Potassium bromate and hydrogen peroxide induced distinct changes in the cellular concentrations of the metabolites glutathione, methionine, N-acetylputrescine, 4-aminobutyrate, and uracil. Symbols and error bars represent median +/− 95% interval of confidence limits of n= 3 and n=4 independent experiments for potassium bromate and hydrogen peroxide, respectively. Metabolites with significantly different relative concentrations are surrounded by a dashed red line. Figure showing metabolomic changes in donor bacterial cells exposed to potassium bromate or hydrogen peroxide. Panels A and B present principal component analysis (PCA) plots of NMR-detected metabolites from donor populations exposed to potassium bromate (1.5 mM) or hydrogen peroxide (10 mM), respectively, compared with mock-treated controls. In both cases, exposed and control populations form distinct clusters, indicating significant differences in metabolic profiles. Panel C compares metabolic profiles between potassium bromate- and hydrogen peroxide-exposed populations and shows that the two oxidative agents induce distinct metabolic states. Each dot represents one biological replicate from three or four independent experiments, and coloured ovals indicate 95% confidence intervals. Panel D shows changes in cellular concentrations of selected metabolites, including glutathione, methionine, N-acetylputrescine, 4-aminobutyrate, and uracil, demonstrating that potassium bromate and hydrogen peroxide produce distinct metabolic responses. Symbols represent medians and error bars indicate 95% confidence intervals. Metabolites with significantly different relative concentrations are surrounded by a dashed red line.

Altogether, these results indicate that, even though similar changes in intracellular levels of several metabolites reflect common cellular responses to both redox stress agents, potassium bromate and hydrogen peroxide differentially affect cellular metabolism during exposure, which may underlie the observed differences in mutagenesis.

## DISCUSSION

### Weak redox stress agents promote mutagenesis during bacterial conjugation and generate distinct mutational spectra

In the present study, we have shown that bacterial conjugation is a process remarkably sensitive to the effect of environmental mutagens. Our results (Fig. 2, 4, and S1) indicate that during conjugation most of the induced mutations arise from damage to the F episome ssDNA T-strand when it is being transferred from the donor to the recipient cell. This finding implies that bacterial conjugation can serve as a system to detect and characterize mutagenesis induced by weak mutagens. Here, as proof of principle, we have employed two environmentally relevant redox stress agents, potassium bromate and hydrogen peroxide.

Analysis of the mutational spectra induced by these compounds at equitoxic concentrations during conjugation shows that they have very distinct effects on F DNA mutagenesis, resulting in overall different changes in the mutational spectra (Fig. 3 and S4). While both redox stress agents increased all types of mutations, hydrogen peroxide preferentially induced base substitutions, specifically at C and at G, while potassium bromate preferentially induced changes at C, mainly +/− 1 BP mutations (Fig. 3). These results are aligned with those obtained from the analysis and comparison of the NMR-detected cellular metabolic landscapes that equitoxic concentrations of potassium bromate and hydrogen peroxide induced in donor populations, which showed significant differences in the levels of various metabolites involved in ameliorating the impact of redox stress (Fig. 5).

Altogether, these results indicate that different redox stress agents exert distinct mechanisms of cellular damage that lead to specific mutational spectra. Nuanced characterization of environmentally relevant redox stress agents may improve our understanding of their roles in the emergence and progression of diverse pathological processes. The conjugation-based system described here, in which ssDNA is transiently exposed, and the resulting damage can subsequently be analysed, represents a powerful approach for mutagenesis characterization. Furthermore, the sensitivity of this system could be enhanced in future studies by employing donor and/or recipient strains carrying mutations that increase their susceptibility to redox stress-induced damage.

### Redox stress agents differentially impact cellular metabolism and induce distinct mutations in bacterial and yeast ssDNA reporter systems

In previous studies employing a ssDNA reporter system in *Saccharomyces cerevisiae*, in addition to identifying the base most vulnerable to mutagenic change, we were able to characterize the mutational motif (the DNA context adjacent to the most frequently mutated base) of potassium bromate, hydrogen peroxide, and paraquat that results from exposure to these oxidizing agents (50,51). These mutational motifs (the DNA context adjacent to the most frequently mutated nucleotide) of each of these compounds are similar, suggesting that specific DNA sequences are more sensitive to oxidative damage. Yet, the preferentially damaged base and resultant mutation were different (G to T for potassium bromate and C to T for hydrogen peroxide and paraquat), indicating a distinct, specific mechanism of mutagenesis for each of these redox stress agents in yeast as well as in bacteria.

A comparison of the consequences of redox stress-induced mutagenesis during conjugation in bacteria with those observed in the yeast ssDNA reporter system underscores their similarity, manifested by the extreme vulnerability of ssDNA to mutagenesis. However, the mutational footprints of potassium bromate and hydrogen peroxide are clearly different between these systems (Supplementary Table S2). Also, contrary to the widely accepted notion of guanine being the most vulnerable target of redox stress mutagenesis (66,67), we found that this is not the case for bacteria during conjugation (Fig. 3). These differences may result from several non-mutually exclusive causes, some of which we can easily ascertain.

In yeast and in other eukaryotes the presence of ssDNA is relatively temporary (e.g., during Okazaki fragments replication or re-synthesis of pre-telomeric regions) or the consequence of molecular malfunctioning (e.g., such as replication fork uncoupling during replication stress in cancers). However, conjugational transfer of ssDNA in bacteria is conserved in many species. Therefore, it is reasonable to assume that multiple bacteria-specific mechanisms have evolved to protect the ssDNA during the transfer. Identification of such protection mechanisms could be important for practical goals of preventing the acquisition of virulence and antibiotic resistance or, vice versa, exaggeration of the damage and blockage of horizontal gene transfer.

In this regard, one of the important differential consequences of exposure to redox stress agents in yeast and bacteria was the unequal cytotoxicity and mutagenicity produced by the same dose of damaging agents. To obtain a significant induction of mutagenesis in the bacterial system that was similar to the mutagenesis generated in the yeast system, we had to employ doses of the mutagens that caused more extensive cell killing (about 85% in bacteria compared to 15 % in yeast). This indicates that the yeast system is more sensitive, i.e., more susceptible to redox stress-induced damage, compared to the bacterial system.

An additional factor to consider is that the reporter genes of both systems are different in DNA context and length. The yeast ssDNA reporter system has been designed for detecting mutations that result in inactivation of the closely spaced *CAN1* and *ADE2* genes. Mutagenesis analysis, using this reporter, results from sequencing these two genes as well as the adjacent *URA3* and *LYS2* genes of *CAN1* and *ADE2* double-mutant isolates. This reporter system thus allows the selection of two or more mutations (68), whereas in the bacterial system developed in our study, a second mutation arising during conjugation would be detected only incidentally. In the yeast system, a 46-fold increase in clustered mutations was reported when MMS-exposed cells were compared to unexposed ones, with clusters comprising a range of 2 to 30 mutations (68). We wanted to investigate if similar MMS-induced clusters are also generated in the F episome during conjugation. We performed whole genome sequencing (WGS) on a random sample of 10 *lacI* transconjugants from an MMS-exposed culture and 10 *lacI* transconjugants from the corresponding mock culture (all from the series of conjugation experiments in minimal media with MMS). The results showed that four out of the 10 MMS-treated *lacI* mutants sequenced carry an additional mutation in F, while none of the mock derived mutants carry a secondary mutation (Supplementary Table S3). These results indicate that multiple mutations occur in ssDNA the bacterial system too. Introducing a second mutational reporter into the F episome, as in the yeast system, would likely increase the frequency of multiple mutations detection and thereby facilitate their analysis in the bacterial system.

Another reason for the differences observed could be that yeast and bacteria have distinct metabolic responses to redox stress. To obtain insight on this issue, we determined the fold change in metabolite levels (relative to mock exposures) determined by NMR spectroscopy and compared the resultant data between these two systems for the same type of exposure (i.e., potassium bromate or hydrogen peroxide). Different groups of metabolites were detected for both systems. For those metabolites that were shared between yeast and bacteria (Supplementary Fig. S6), a subset shows similar behaviour in both systems, notably glutathione in cells exposed to potassium bromate, but others revealed a very different response, such as ADP and succinate in cells exposed to any of the two oxidants. This suggests that the metabolic pathways responding to these redox stress agents may differ substantially between the two systems.

In summary, the damaging effects of redox stress agents on DNA and other cellular components appear to depend not only on the specific agent, as already shown (50,51), but also in the specific cellular system exposed to that agent. This suggests that the overall cellular effects from exposure to redox-stress agents might be tissue-dependent. Studies published on potassium bromate-induced mutagenesis appear to support such conjecture as this redox agent induced distinct mutational spectra depending on the cellular context: large deletions in lymphoblastoid TK6 cells (69); increased A:T to T:A base substitutions and single-base deletions in kidneys of rats (70); and C:G to A:T base substitutions, as well as base pair insertions and deletions, in human epithelial cells (71).

### Bacterial horizontal gene transfer is highly sensitive to mutagens

Our finding that conjugation is highly sensitive to environmental mutagens has profound implications on bacterial evolution and adaptation to antibiotics. Conjugal plasmids encode many different genes that promote not only their own survival, but commonly also the survival of their bacterial host too, such as virulence and antibiotic resistance genes. In this sense, de novo acquisition of antibiotic resistance most frequently takes place through conjugation (reviewed in (37,72)). In natural environments, bacteria are continuously exposed to redox stress agents (reviewed in (73)). For example, pathogenic bacteria are exposed to hydrogen peroxide and other ROS released by the host immune cells during infection (reviewed in (47)) or the commensal bacteria that comprise a large component of the human gut microbiome may be exposed to oxidizing agents such as potassium bromate, present in food (reviewed in (74)). Our findings suggest that these agents may induce mutations even at low exposure doses in conjugative plasmids during ssDNA transfer from donor to recipient, potentially enhancing bacterial adaptation as well as increasing virulence or antibiotic resistance. It is noteworthy that these agents only moderately reduce conjugation efficiency, even at concentrations that result in approximately 85% cell death (Supplementary Fig. S7), which indicates that conjugation has evolved into a highly resilient process that helps ensure the survival of the transferred plasmid.

Conjugation has traditionally been regarded primarily as a mechanism of horizontal gene transfer through which bacteria acquire new genetic traits. Our data supports the concept that it is also a relevant process to generate genetic diversity. First, the fact that bacterial cells carry multiple copies of the same conjugative plasmid favours the accumulation of recessive mutations and the generation of novel alleles in some copies that can be expressed phenotypically after episome segregation during cell division or after conjugation (75,76). The results from our experiments using *lacZ* as reporter (Fig. 4) suggest that mutations induced in *lacI* in donor cells (Supplementary Fig. S5) were not detected most probably because they were recessive. Yet, these mutations can potentially be transferred in ulterior conjugation events. Second, the fact that conjugation per se is mutagenic for the conjugative plasmid being transferred ((39,53), and this study). And third, as we also show here, the fact that the ssDNA transferred during conjugation is extremely sensitive to the effect of environmental growth conditions (media) and mutagens.

In future studies it will be important to determine if this distinctive mutational sensitivity is the result of selection or simply an unavoidable consequence of the conjugative process in bacteria and plasmids. In any case, this characteristic likely contributes to the adaptation of pathogenic bacteria to host environments and antibiotic exposure and provides an additional rationale for strongly pursuing strategies to limit bacterial conjugation, especially in environments conducive for the transfer of virulence factors and antibiotic resistance (77,78).

## ACKNOWLEDGEMENTS

We thank Rajula E. Alleva, Joanna B. Goldberg, and Roel Schaaper for their critical review of the manuscript and helpful suggestions. We are grateful to the NIEHS Genomics Core Laboratory for sequencing support, particularly Kevin Gerrish and Leilei Zhang for their technical expertise and assistance. We thank Ashley M. Brooks for assistance with data analysis, and Ying Liu and Marcus Jackson for assistance with data and file management.

## AUTHOR CONTRIBUTIONS

Libertad García-Villada: Methodology, Investigation, Formal analysis, Writing—original draft. Brett A. Shore: Investigation, Writing—review & editing. Kasey Kiser: Investigation. Isabella Russ: Investigation. Scott A. Gabel: Methodology, Validation, Investigation. Geoffrey A. Mueller: Methodology, Validation, Writing—review & editing. Natalya P. Degtyareva: Conceptualization, Writing—original draft. Paul W. Doetsch: Resources, Writing—review & editing.

## SUPPLEMENTARY DATA

## DATA AVAILABILITY

The whole genome sequencing data reported in this study are currently being deposited in NCBI SAR. Accession numbers will be provided during manuscript review and will be made publicly available upon publication. The processed data and analysis code are available on the CEBS database.

