## Supplementary material for "Redox stress agents strongly enhance mutagenesis during horizontal gene transfer in bacteria and leave distinct mutational and metabolic footprints"

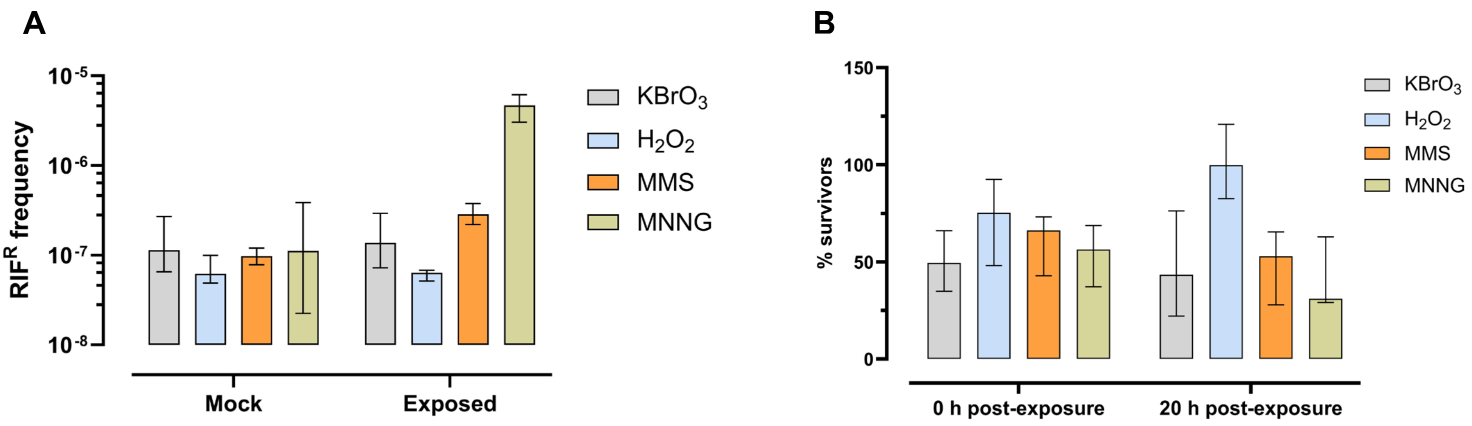


**Supplementary Figure S1.** Exposure to potassium bromate (2.5 mM) or hydrogen peroxide (10 mM) do not induce mutagenesis in chromosomal *rpoB* gene. To verify that potassium bromate and hydrogen peroxide do not induce significant damage to dsDNA under our experimental conditions, the frequency of *rpoB* mutants (CFU of rpoB mutants per ml / total CFU per ml) selected on RIF (100 ug/ml) plates was compared between unexposed ("Mock") and exposed donor populations. Results are shown in (**A**). No significant induction of mutagenesis was observed for potassium bromate or hydrogen peroxide. A small (2.9-fold), but significant (P= 0.0012; paired t test on ln-transformed data) induction was observed for MMS. The alkylating agent methyl nitro nitroso guanidine (MNNG), which is considered a strong dsDNA mutagen, was used as experimental control. As expected, mutagenesis was strongly (42-fold) induced by MNNG. Graph in (**B**) represents the percentage of survivors ((CFU per ml in exposed cultures / CFU per ml in unexposed cultures) x 100) right after exposure and 20 hours after exposure (see Material and Methods). In both figures, bars and error bars represent median +/- 95% interval of confidence limits of n= 4 biological replicates.

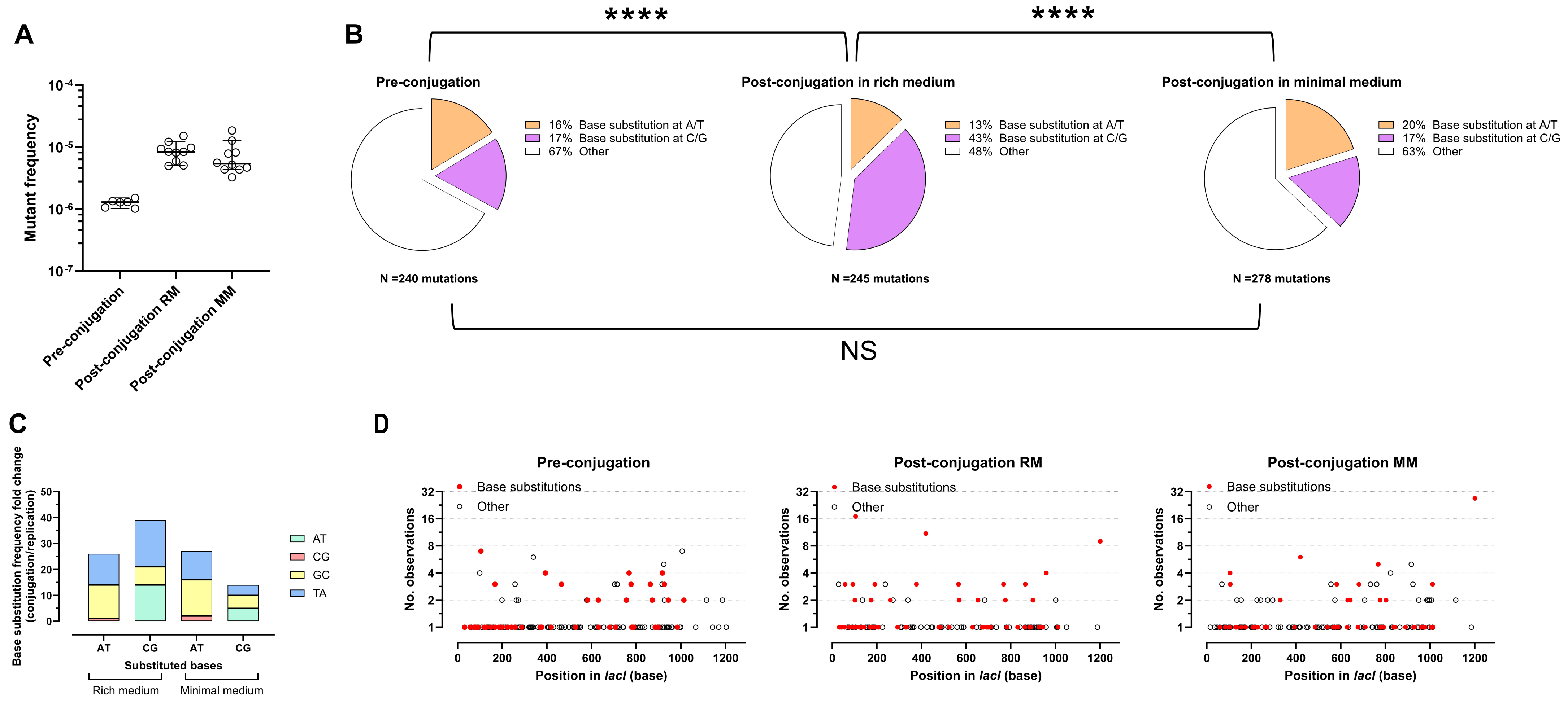


**Supplementary Figure S2.** *lacI* conjugation-induced and replication (vegetative) mutation spectra are qualitatively different. (**A**) The frequency of *lacI* mutants in the transconjugant population ("Post-conjugation" in both, rich and minimal media; RM and MM, respectively) is significantly (~7-fold) higher that in the donor population ("Pre-conjugation"). The horizontal line and error bars represent median +/- 95% interval of confidence limits of n≥ 6 independent experiments. (**B**) Spectra of mutations identified by Sanger sequencing of *lacI* mutants from donor populations ("Pre-conjugation") and transconjugant populations ("Post-conjugation") resulting from conjugation in rich and in minimal medium. Conjugation in rich medium shows an increase in the proportion of base substitutions at C/G that results in a significantly different spectrum of mutations when compared to Pre-conjugation or Conjugation in minimal medium spectra (P < 0.0001 in both comparisons; chi-square test). **(C)** Relative fold change in base substitution frequencies after conjugation in rich and in minimal medium compared to pre-conjugation. (**D**) Distribution of base substitutions (red circles) and other mutations (white circles representing +/- 1base pair changes, >1 base pairs insertions and deletions, duplications, and complex mutations) in *lacI* mutants before conjugation ("Pre-conjugation") and after conjugation in rich and in minimal medium (RM and MM, respectively).

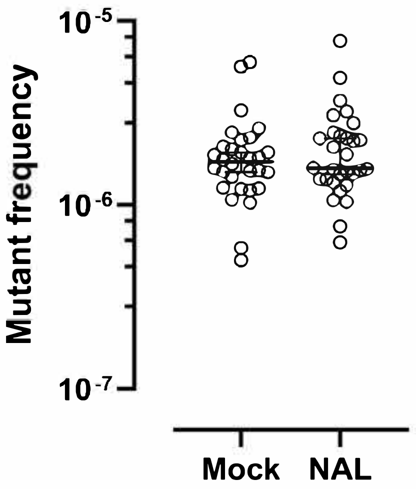


**Supplementary Figure S3.** NAL does not induce lacI mutagenesis on selective plates. Single colonies of WT transconjugants isolated on minimal medium plates supplemented with STR (100 µg/ml) and NAL (40 µg/ml) were grown ON in LB supplemented with antibiotics (NAL and STR) and a sample from each culture was plated on Pgal medium with or without NAL (40 µg/ml). No increased mutagenesis was observed on the samples plated on NAL. The horizontal line and error bars represent median +/- 95% interval of confidence limits of n≥ 10 biological replicates from each of three independent experiments.

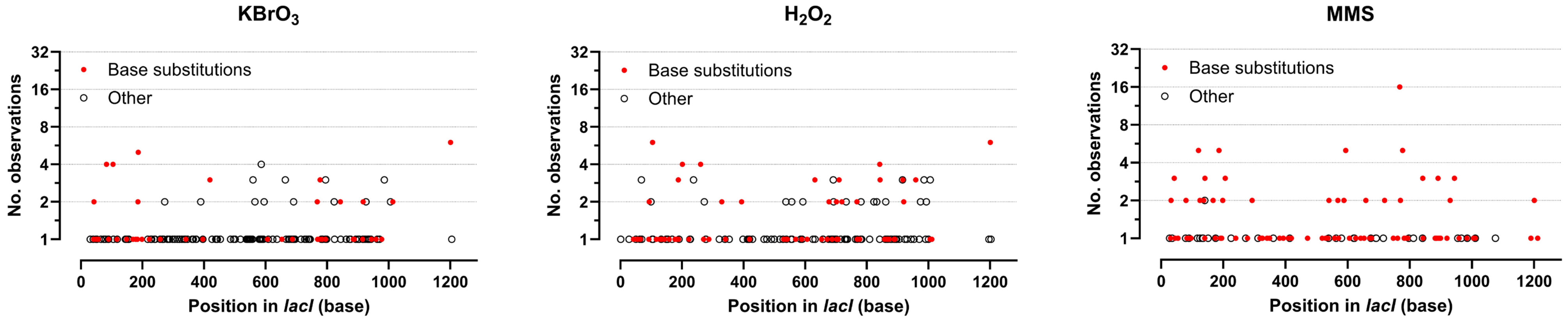


**Supplementary Figure S4.** Differences in mutational spectra among mutagen-exposed populations is not due to hot spots. Distribution of base substitutions (red circles) and other mutations (white circles representing +/- 1base pair changes, >1 base pairs insertions and deletions, duplications, and complex mutations) in lacI mutants from potassium bromate-, hydrogen peroxide-, and MMS-exposed transconjugant populations.

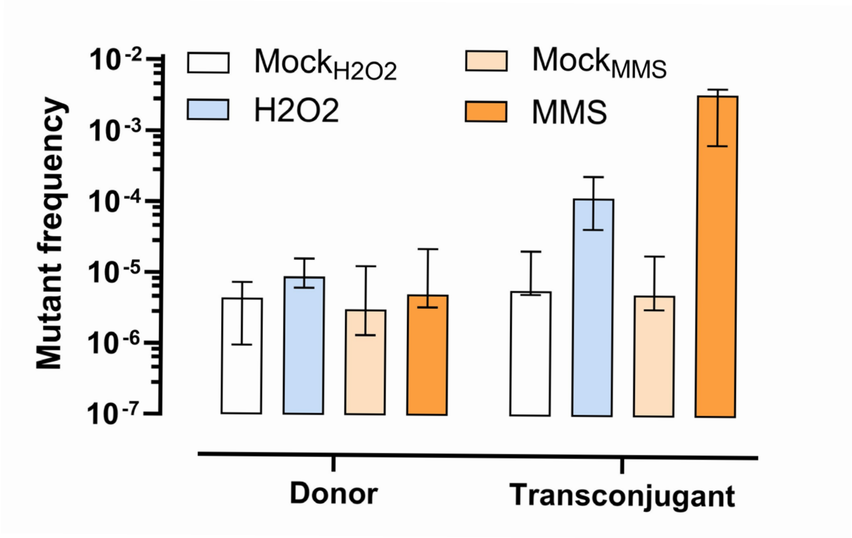


**Supplementary Figure S5.** No significant increase in *lacI* mutant frequency was observed in the donor population after conjugation in the presence of hydrogen peroxide or MMS. Exposure to hydrogen peroxide (10 mM) or to MMS (10 mM) during conjugation did not increase the *lacI* mutant frequency (CFU of lacI mutants per ml / CFU of survivors per ml) in donor populations. Bars and error bars represent median +/- 95% interval of confidence of n≥ 4 independent experiments.

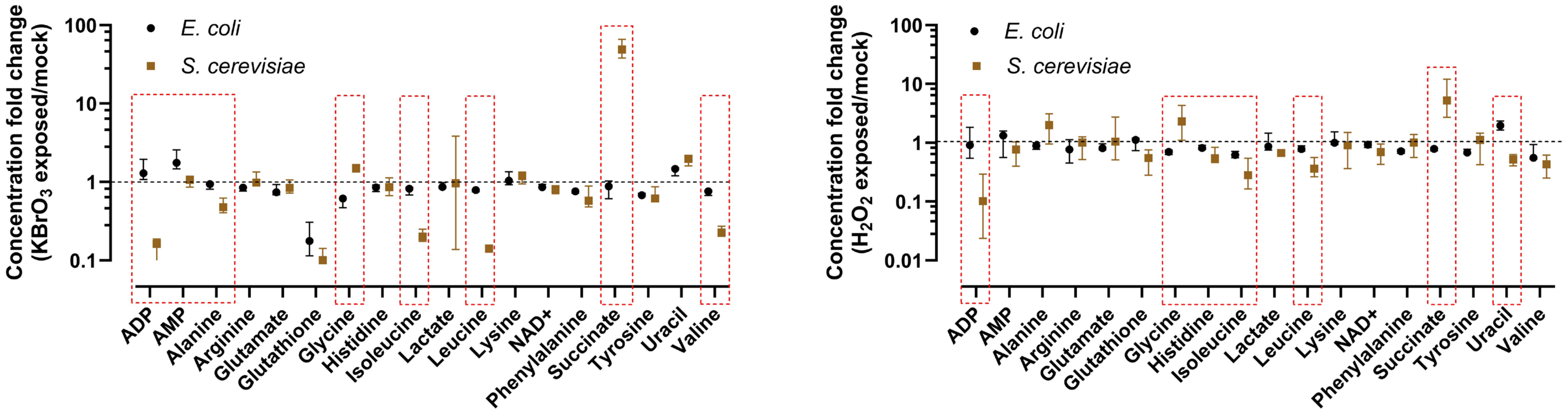


**Supplementary Figure S6.** Metabolic landscape differences between *E. coli* and *S. cerevisiae* exposed to potassium bromate and hydrogen peroxide. Graphs show the relative (with respect to mock) fold change in the cellular concentration of those metabolites (18) detected through $^{1}$H-NMR spectroscopy common for both organisms. *S. cerevisiae* data is from (Degtyareva et al., 2023). Symbols and error bars represent median +/- 95% interval of confidence limits of n≥ 3 independent experiments. Metabolites with significantly different relative concentrations in both systems are surrounded by a dashed red line.

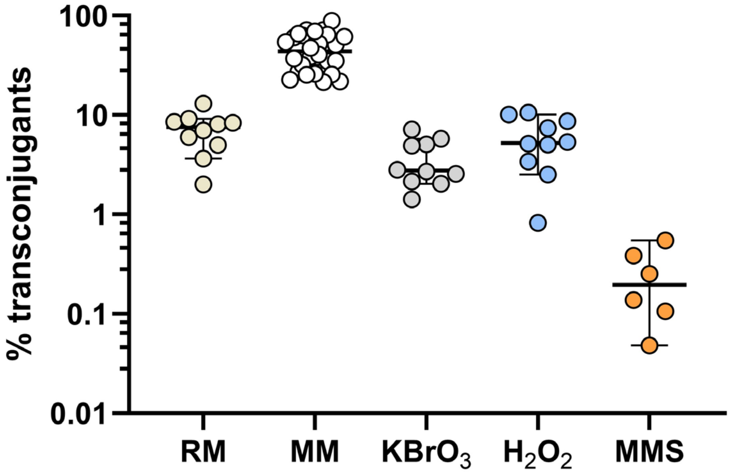


**Supplementary Figure S7.** Conjugation efficiency is only moderately reduced by oxidizing agents. The graph represents the percentage of transconjugants (CFU transconjugant cells per ml / CFU recipient cells per ml) resulting from conjugation in rich medium (RM) and in minimal medium containing none (MM) or different mutagens: potassium bromate (1.5 mM), hydrogen peroxide (10 mM), and MMS (10 mM). Symbols and error bars represent median +/- 95% interval of confidence limits of n≥ 6 independent experiments.

**Supplementary Table S1.** Distribution of +/- 1 BP with respect to homopolymeric nucleotide runs.

| Run size | Pre-conjugation | Mock RM[1] | Mock MM[2] | $KBrO_3$ | $H_2O_2$ | MMS |
|---|---|---|---|---|---|---|
| **0** | 0 | 1 | 3 | 1 | 0 | 0 |
| **1** | 14 | 7 | 27 | 34 | 27 | 11 |
| **2** | 9 | 8 | 11 | 18 | 14 | 3 |
| **3** | 5 | 7 | 5 | 19 | 12 | 4 |
| **4** | 6 | 4 | 6 | 2 | 3 | 1 |
| **5** | 8 | 3 | 2 | 2 | 3 | 1 |
| **Total** | **42** | **30** | **54** | **76** | **59** | **20** |

[1] RM, Rich medium

[2] MM, Minimal medium

**Supplementary Table S2.** Bacterial and yeast ssDNA oxidative damage reporters render different results.

| **Type of exposure** | **Prevalent damaged base** | |
|---|---|---|
| | **In *E. coli*** | **In *S. cerevisiae*** |
| **Mock MM[1]** | C (34%) and G (28%)* | G (53%) |
| **$KBrO_3$** | C (32%) | G (70%) |
| **$H_2O_2$** | C (41%) and G (43%) | C (55%) |
| **MMS** | C (41%) | C (%75) |

[1] MM, Minimal medium

**Supplementary Table S3.** F episome mutations detected in *lacI* mutants isolated from mock- and MMS (10 mM)-exposed transconjugant cultures.

| Exposure | Mutant no. | Region | Type | Reference | Allele | Length | Gene |
|---|---|---|---|---|---|---|---|
| Mock | 1 | 222091..222547 | Deletion | | | 462 | *lacI* |
| | 2 | 222693^222702 | Duplication | | CCAGGATGCC | 10 | *lacI* |
| | 3 | 222357^222358 | Duplication | | TGC | 3 | *lacI* |
| | 4 | 222734^222735 | Insertion | | C | 1 | *lacI* |
| | 5 | 222901..2229019 | Deletion | | | 25 | *lacI* |
| | 6 | 222492 | SNV | T | A | | *lacI* |
| | 7 | 221919 | SNV | T | C | | *lacI* |
| | 8 | 222734^222735 | Insertion | | C | 1 | *lacI* |
| | 9 | 223015^223016 | Insertion | | GAACC | 5 | *lacI* |
| | 10 | 222109 | SNV | C | T | | *lacI* |
| MMS | 1 | 222352 | SNV | A | T | | *lacI* |
| | 2 | 213135 | SNV | C | A | | |
| | 2 | 222526 | SNV | T | A | | *lacI* |
| | 3 | 224676 | Deletion | G | - | 1 | *mhpA* |
| | 3 | 223078 | SNV | C | T | | *lacI* |
| | 4 | | No mutation | | | | |
| | 5 | 213936 | SNV | T | G | | |
| | 5 | 222278 | SNV | G | T | | *lacI* |
| | 6 | | No mutation | | | | |
| | 7 | 172752 | Deletion | T | | | |
| | 7 | 222979 | SNV | T | A | | *lacI* |
| | 8 | 222226 | SNV | T | G | | *lacI* |
| | 9 | 222352 | SNV | A | T | | *lacI* |
| | 10 | 222350 | SNV | C | T | | *lacI* |

SNV: Single Nucleotide Variant
